# A genomic link in China roses: and they all lived prickly but water deficient ever after?

**DOI:** 10.1101/2020.07.16.207795

**Authors:** Mi-Cai Zhong, Xiao-Dong Jiang, Guo-Qian Yang, Wei-Hua Cui, Zhi-Quan Suo, Wei-Jia Wang, Yi-Bo Sun, Dan Wang, Xin-Chao Cheng, Xu-Ming Li, Xue Dong, Kai-Xue Tang, De-Zhu Li, Jin-Yong Hu

## Abstract

Prickles act against herbivores, pathogens or mechanical injury, while also prevent water loss. However, whether prickles have new function and the molecular genetics of prickle patterning remains poorly explored. Here, we generated a high-quality reference genome assembly for ‘Basye’s Thornless’ (BT), a prickle-free cultivar of *Rosa wichuraiana,* to identify genetic elements related to stem prickle development. The BT genome harbors a high level of sequence diversity in itself and between cultivar ‘Old Blush’ (*R. chinensis*), a founder genotype in rose domestication. Inheritance of stem prickle density was determined and two QTL were identified. Differentially expressed genes in QTL were involved in water-related functions, suggesting that prickle density may hitchhike with adaptations to moist environments. While the prickle-related gene-regulatory-network (GRN) was highly conserved, the expression variation of key candidate genes was associated with prickle density. Our study provides fundamental resources and insights for genome evolution in the Rosaceae. Ongoing efforts on identification of the molecular bases for key rose traits may lead to the improvement of horticultural markets.

## Introduction

Understanding the molecular genetic mechanisms driving morphological variation remains a great challenge. When we say, “there are no roses without thorns” we employ a metaphor meaning we are willing to endure something unpleasant because it is associated with something good. In fact, rose prickles are acuminate protuberances formed by the deformation of plant trichomes that are often mistaken for true spines or thorns. Prickles are usually interpreted as epidermal adaptations protecting plants from herbivores, pathogens or mechanical injury. In the children’s book, *Wild Animals I have Known* (1898), author Ernest S. Thompson depicted the rose bush as a protector of cottontail rabbits while the same prickles harmed people and other animals. However, prickles also increase the thickness of epidermis which reduces heat and water dissipation [1, 2]. Many plants in the families Rosaceae, Araliaceae, Fabaceae and Rutaceae bear prickles on their leaves, stems, and fruits. Trichomes still serve as developmental models to study the control of cell proliferation and growth in *Arabidopsis and Gossypium* thus allowing the patterning gene-regulatory-network (GRN) to be investigated in detail [1, 3–6]. However, the molecular and genetic mechanisms underlying stem prickle development have not been studied systematically.

With more than 36,000 cultivars, roses have long been revered for their beauty and fragrance, despite the presence of abrasive prickles on their stems. Roses offer far more morphological novelty, including continuous flowering (CF), prickles, multi-petal corollas and complicated floral scents absent in such current model systems as *Arabidopsis*, rice and poplar. Therefore, rose serves as an additional woody plant model for studying important traits with commercial values [7, 8]. However, rose genetics are unusually complex due to past histories of natural inter- and intra-specific hybridization, polyploidy, and unbalanced meiosis further exaggerated under human selection [9–11]. Determination of a highly heterozygous genome for roses remains challenging, as seen by the very fragmented assembly trials on three species [12, 13]. Currently two versions of chromosome-level genome sequences for doubled-haploid materials of one cultivated founder genotype (*Rosa chinensis* ‘Old Blush’, haploOB) have been described [12, 14]. Enormous sequence variations can exist between rose cultivars as in maize and they may contribute to key agricultural traits [15]. Therefore, a complete picture of genome assembly for diploid roses is necessary for comparative genomics to identify the molecular genetic bases of key morphological traits pursued after centuries of breeding.

The glandular-trichome-based prickles on rose stems [16] are inconvenient in floriculture and commercial nurseries. Although most rose genotypes harbor prickles the “prickle-free” cultivar known as ‘Basye’s Thornless’ (BT) is a variant of *R. wichuraiana* Crep., a diploid founder species (2n=14) in rose domestication (Fig. 1a) [8, 17–19]. It has been used to create hundreds of commercial hybrids with ‘Pink Roamer’ as the first [18]. BT grows in a relatively moist environment. The BT cultivar has at least six traits differing from the OB cultivar, a founder in hybridization used even more frequently. Compared to OB, the BT flowers only once a year, bears prickle-free stems and shows prostrate growth. While each flower bears only a single whorl of white petals it continues to show a higher resistance to black spot [18]. This explains why BT is often crossed with OB. Unfortunately, molecular information on BT is limited [14, 20–26]. Here, we determined the chromosome-level genome sequences of BT. We systematically analyzed the prickle QTL and GRN. We hypothesize that prickle-free may hitchhike on genomic segments selected for adaptation to relatively warm and humid climates.

**Fig. 1.**
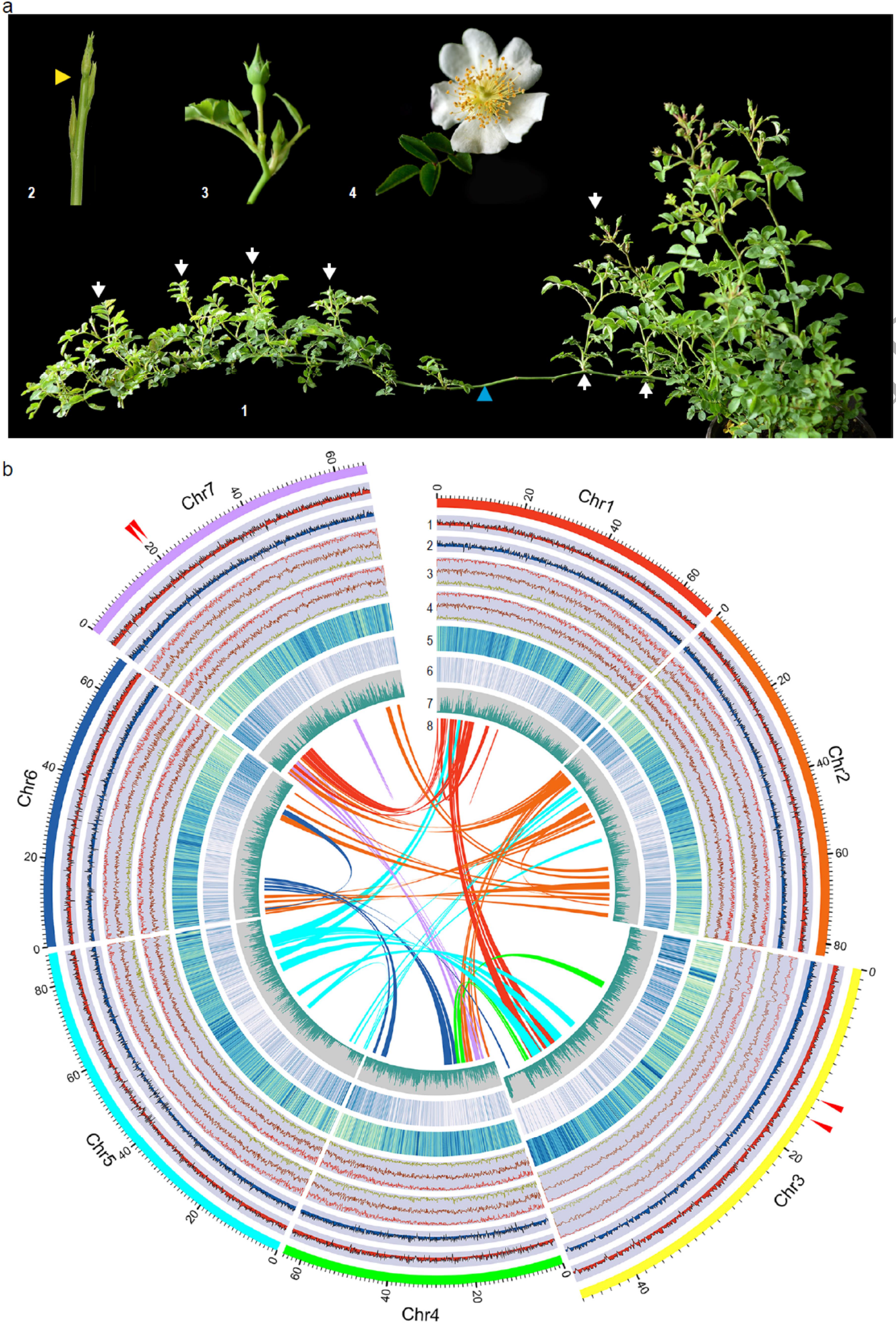
Anatomical and genome features of *R. wichuraiana* ‘Basye’s Thornless’ (BT). **a.** Growth habit of flowering BT plant in April, 2020. **1**, Branches bearing flowering buds erect/sub-erect (white arrows) while vegetative stems remain prostrate (blue arrowhead). **2**, Early flowering stage (orange arrowhead). **3**, Flower bud prior to anthesis. **4**, Fully open flower. **b.** Landscape of genome features for BT genome. **1**, Gene expression profiles for leaf materials grown in March 2017. **2**, Gene expression profiles for leaf materials grown in November 2016. **3** and **4**, DNA methylation patterns for samples in **1**(**3**) and **2**(**4**). Lines in red, dark red, and brown mark the CG, CHG, and CHH methylation, respectively. **5**, Contents of transposable elements (TEs) in each chromosome. **6**, Gene density in each chromosome. **7**, SNP density along chromosomes in every 100kb bins. **8**, internal syntenic blocks within BT chromosomes. Arrow heads marked the two QTL regions.

## Results and discussion

### *De novo* chromosome-level assembly of the BT genome

Flow cytometry showed that the genome size of BT was estimated at 525.9 (±5.91) Mb. The initial genome survey analysis with ~267x Illumina reads predicted a genome size of 525.5 Mb with ~1.03% heterozygosity. We generated ~93x PacBio Sequel and ~90x Oxford Nanopore Technology (ONT) reads and used a *CANU* based hybrid strategy to construct contigs (Fig. S1; Table S1, S2). We obtained 1,554 contigs of 530 Mb (Table S3, S4). We built a chromosome-level assembly using 34.64 million unique Di-Tags read pairs produced via Hi-C experiment (Table S5). Hi-C analysis generated 2,711 contigs, among which 857 (52.97% in 1,618 clustered) contigs were ordered and oriented into seven pseudo-molecules spanning a 481.77 Mb (92.41% of 521.32 Mb total sequences clustered; Table S6). There were 46.18 Mb sequences not ordered/oriented. After manual correction, we obtained the final assembly of the BT genome (530.07 Mb). Chr5 was the longest (87.44 Mb) while Chr3 the shortest (46.04 Mb) (Fig. 1b; Fig. S2; Table S6, S7).

### Reference-quality and high completeness of the BT genome

We used several methods to evaluate the quality and completeness of the BT genome. First, this genome was subjected to Benchmarking Universal Single-Copy Orthologs (BUSCO, embryophyta_odb10) analysis. Overall, 93.9% (2,184; complete) and 3.0% (69; fragmented) of the 2,326 expected plant gene models were identified (Table S8). Two independent genetic maps based on different populations were used to assess the quality of the BT genome. The collinearity with the K5 genetic map [27] was very high with most of the association efficiency above 95% (Table S9; Fig. S3). Chr4 was an exception in the OBxBT genetic map, in which the distribution of molecular markers was obviously skewed by high proportion of distorted markers [25]. The remaining six chromosomes showed collinearity at ~90% correlation rate (Table S9; Fig. S4). We next aligned our BT assembly to the recently reported haploOB genome assemblies [12, 14], and identified a very high collinearity (Fig. 2a, 2b; Fig. S3). This revealed a high synteny level and gene order between the genotypes. We re-mapped and obtained high mapping rates of the Illumina (98.16%) and PacBio (96.39%) reads (Table S10). An per-base estimate of the quality-value (QV) reached 34.61. Finally, the genome-wide LTR Assembly Index value of BT was 20.04, a level of golden reference (Fig. S5). Collectively, these indicated the BT genome was in high-quality and completeness comparable to the two haploOB genomes (Table S11).

**Fig. 2.**
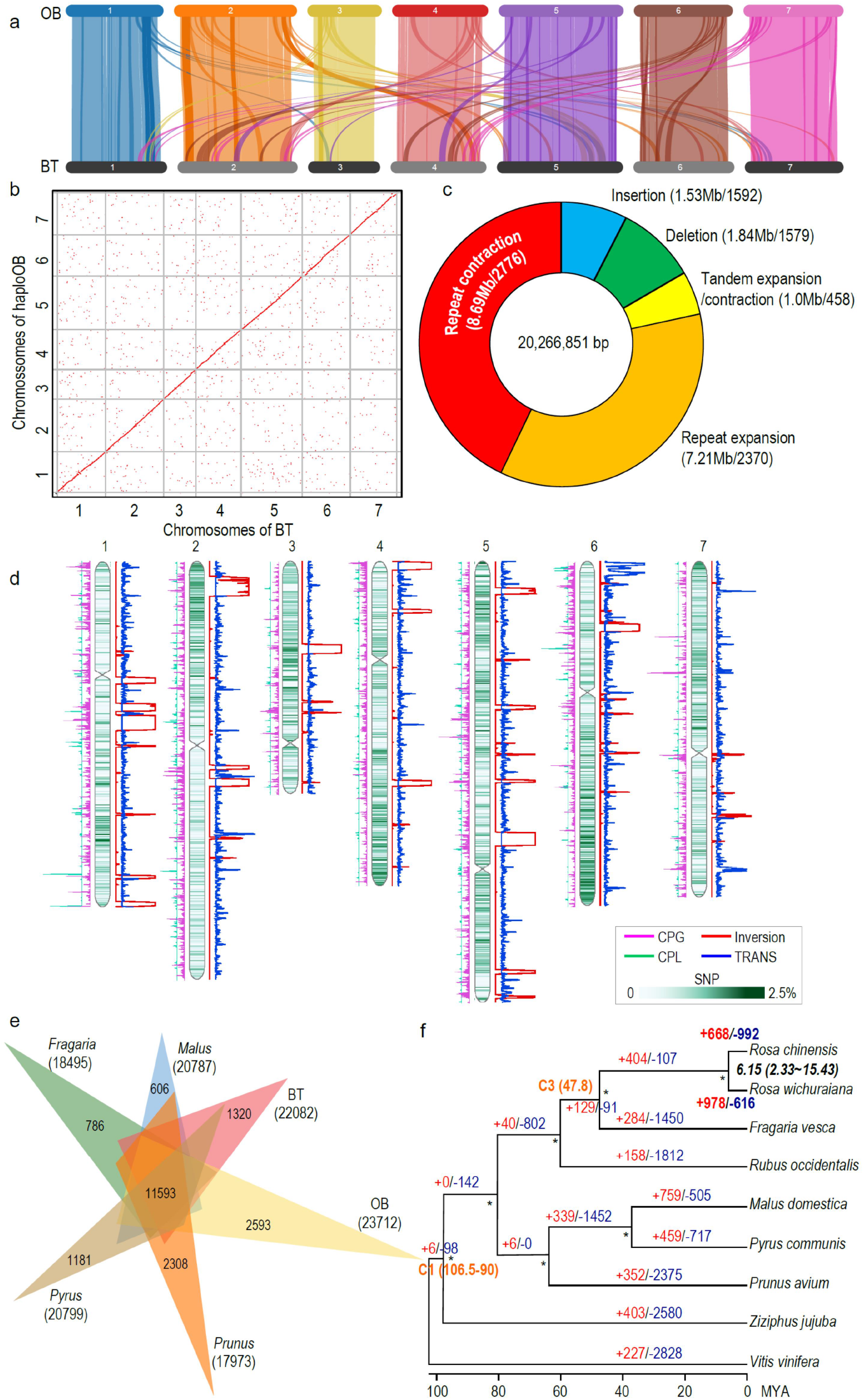
Comparative genomic analyses between BT, OB and other members of the Rosaceae. **a.** Gene collinearity (minimum ten syntenic gene pairs) between the BT and Raymond’s haploOB genomes. High level of collinearity between chromosome pairs indicated by light color lines/blocks. Dark color lines mark genome segments with relatively lower collinearity. **b.** Macrosynteny patterns between BT and haploOB (Raymond et al. 2018) genomes along each chromosome. Dots closest to the diagonal line represent collinearity between the two genomes with <10 Kb fragments filtered out. **c.** Doughnut chart showing large-size sequence variation between BT and haploOB. Numbers in brackets indicate the total length of each type of variation (before /) and the number of events (after /). **d.** Genome alignment between BT and haploOB identifying significant structural variants including genome inversions (red lines), translocation (TRANS; blue lines), copy gain (CPG; cyan lines; in OB with BT as reference) and copy loss (CPL; green lines). Data plotted as per type of variation in 1 Mb of genomic region with step size of 100 kb by length density. SNP density plotted on the chromosomes with 2.5 SNPs per 100 bp length at the highest level. **e.** Comparison of the numbers of gene families identified via OrthoMCL. Venn diagram showing shared homologous gene families among genomes of BT, haploOB, *Fragraria*, *Malus*, *Pyrus* and *Prunus*. Numbers in brackets indicate total gene families. Only numbers of genome-specific and shared among six genomes shown. Numbers in brackets show total gene families for each species. **f.** Phylogenetic relationships of BT with other taxa in the Rosaceae based on 1,230 shared single-copy orthologous genes with *Ziziphus jujuba* and *Vitis vinifera* (outgroups). Estimated divergence time between BT and OB following the fossil calibrations of nodes C1 (106.5-90 MYA) and C3 (47.8 MYA) (in orange, according to Zhang et al. 2018). Numbers on branches indicate the counts of gene family expansion (+, in red) and contraction (-, in blue) along each lineage. * gives the bootstrap supports above 95% in 500 times simulation.

### A high proportion of *Copia* type LTRs in the BT genome

Repeat sequences represented about 65.19% (345.58Mb) of the BT genome, in which LTR retrotransposons were about 45.09% (or 69.16% of all repeats; Fig. 1b; Table S12). In contrast to haploOB [12, 14], but similar to the *Musa balbisiana* assembly [28], the BT genome had much more *Copia* elements (28.29% of the genome, or 62.74% of all LTRs) than *Gypsy* (16.39%). The distribution of *Copia* type LTRs was antagonistic to that of the *Gypsy* type on almost all BT chromosomes (Fig. S6). The ~0.58 *Gypsy*-to-*Copia* ratio was one of the lowest in the sequenced species. It suggests that *Copia* amplification contributes much more than other types of TEs to BT genome evolution (Table S13; Fig. S7). In general, the dense distribution of TEs in the pericentromeric regions correlated inversely with gene density on the BT chromosomes (Fig. 1b).

### The BT genome has 32,674 protein-coding genes

Gene annotation of the BT genome generated a total of 32,674 protein-coding genes, with an average gene length of ~3.28kb, an average exon length ~251bp and a mean intron length of 390bp (Fig. 1b; Table S14-16; Fig. S8). Protein-coding genes showed focused distribution and high expression levels on chromosome arms, on which highly variable levels of DNA methylation were also observed (Fig. 1b). Furthermore, 15,357 genes were annotated with gene ontology (GO) information (Fig. S9; Table S16). Among these 1,703 genes (~5.21%) encode for transcription factors (TFs) in 69 families while 401 genes (~1.23%) encode for transcriptional regulators in 24 families (Table S17). We predicted 557 tRNAs in 24 families, 78 rRNAs in 4 families, 71 miRNAs in 20 families, and 4,254 pseudo-genes (12,490,896 bp or 2.58%; Table S18).

### The BT genome features a high proportion of allelic sequence polymorphisms

Rose genomes were highly heterozygous [9–14]. Therefore, we next examined sequence polymorphisms within the BT genome. We found 2.3 million SNPs (0.44% of BT genome), in which 1.45 million were transitions and 0.85 million were transversions (Table S19, S20; Fig. S10-S12). About 99.7% of the SNPs were heterozygous with half in intergenic regions. These SNPs resulted in loss/gain of their start or stop codons in 5,204 genes. We further identified 0.51 million indels, in which 520 lead to loss/gain of start/stop codons. Around 44.38% indels were located in intergenic regions (Table S21). These sequence polymorphisms will be valuable in the investigation of allele-specific expression, epigenetic regulation, genome structure and in the evolutionary analyses of roses.

### BT diverges significantly from the haploOB genomes

To understand genetic variation associated with the contrasting traits between BT and OB, we identified 95 synteny blocks covering 87.59% of BT (464.31 Mb) or 80.59% of haploOB (415.5 Mb) genomes. We detected 20 inversion events (covering 1,464 genes of BT or 1,718 genes of haploOB) (Fig. 2a; Table S22, S23). We detected 7.31 million of SNPs and 5.16 millions of one-nucleotide indels as well as 20.27 Mb large structural variants including indels. These included sizes above 50nt (3,171x events) and tandem (458x) or repeat (5,146x) expansion/contractions (Fig. 2c, 2d; Fig. S13; Table S24). We detected 17,034 genomic rearrangement events (~65.8 Mb) not randomly distributed along chromosomes (Fig. 2d; Table S25). Of greater interest, genes related to DNA integration and metabolic processes were significantly enriched in copy-gain-regions (Table S26). This was in line with the strong requirement of maintaining genome stability during frequent hybridization and genomic rearrangement during the long-term domestication of OB [12]. Taken together, a strong sequence divergence occurred between the two genotypes, and this divergence could be used for genetic dissection of important rose traits.

### BT experiences significant gene family expansion/contraction profiles

We identified 11,593 homologous gene families shared among BT, haploOB, and in the genera *Prunus*, *Pyrus*, *Fragaria* and *Malus*. We detected 1,320 gene families with 1,999 genes specific for BT, and found 2,593 families with 4,586 genes unique for haploOB (Fig. 2e). The BT-specific genes were significantly (FDR<0.05) enriched in GO terms for nucleic acid binding, RNA-dependent DNA biosynthesis, and DNA integration processes but proved significantly less for oxidation-reduction genes (Table S27). By comparison, the haploOB-specific genes were functionally over-represented in DNA metabolic, organic substance metabolic, hetero- and organic-cyclic compound binding. These results suggested that BT and OB harbored different and specific gene profiles. However, BT and OB had a very similar within-genome duplication pattern (Fig. S14). Rose and strawberry shared an ancient whole-genome-duplication (WGD) event, but without the recent one featured by apple (Fig. S15, S16) [29].

To understand the phylogenetic position and identify the expanded/contracted gene families in BT we searched for the conserved orthologous gene families in genomes of BT, haploOB, *Malus domestica* Borkh., *Fragaria vesca*, *Prunus avium* L., *Prunus persica* (L.) Batsch., *Pyrus communis* L., and *Rubus occidentalis* L. (all members of the Rosaceae) with *Vitis vinifera* L. (Vitaceae) and *Ziziphus jujube* Mill. (Rhamnaceae) as outgroups. We identified 1,220 single-copy orthologous groups and used them to construct a maximum-likelihood tree (Fig. 2f). BT and OB were sister and *Fragaria* was the closest genus. Following the fossil calibrations of nodes C1 and C3 for the Rosaceae [30], the estimated divergence time of BT and OB was ~6.15 MYA (95% highest posterior density: 2.33−15.43 MYA). A total of 978 and 616 gene families were apparently expanded and contracted, respectively, in the BT genome (Fig. 2f). This was significantly different from that of OB (*Yates’ Chi-square test*, p<0.0001; Fig. S17; Table S28). Being consistent with its high resistance to black spot [18], BT expanded significantly the *NAC* family genes (146 against 114 in OB; p=0.05; Table S17) especially on clade V (72 in BT and 41 in haploOB, p=0.004; Fig. S18). The lineage-specific expansion and expression divergence of *FAR1/FRS-like* genes correlated well with transition of shoot growth behaviors upon flowering in BT [31]. These corroborated with the fact that BT and haploOB had a high level of sequence divergence and morphological variation due to the time of divergence between them (Fig. 2a-2d).

### Genetic patterning of stem prickle density in BTxOB populations

We scored stem prickle density in the 99-individual F1 and the 148-individual BC1F1 populations. Both were generated via crossing the prickle-free BT and prickly OB (Fig. 3a; Fig. S19-S21) [22, 25]. In F1s, the number of prickle-free vs. prickly individuals was 13:86, a ratio not deviating from 1:7 (three loci hypothesis, *Chi-square test*, p=0.96) or 1:15 (four loci, p=0.43). The BC1F1 plants, based on back-crossing one prickle-free F1 individual to a prickly OB, showed a normal distribution of prickle density (*Kolmogorov-Smirnov normality test*, D = 0.043, p = 0.723; Fig. S22). Ten plants were prickle-free and the remaining 138 had prickles (Fig. 3b, 3c). Stem prickle density is thus best interpreted as a quantitative trait regulated by 3-4 loci with the being prickly as incomplete dominant (Fig. S23).

**Fig. 3.**
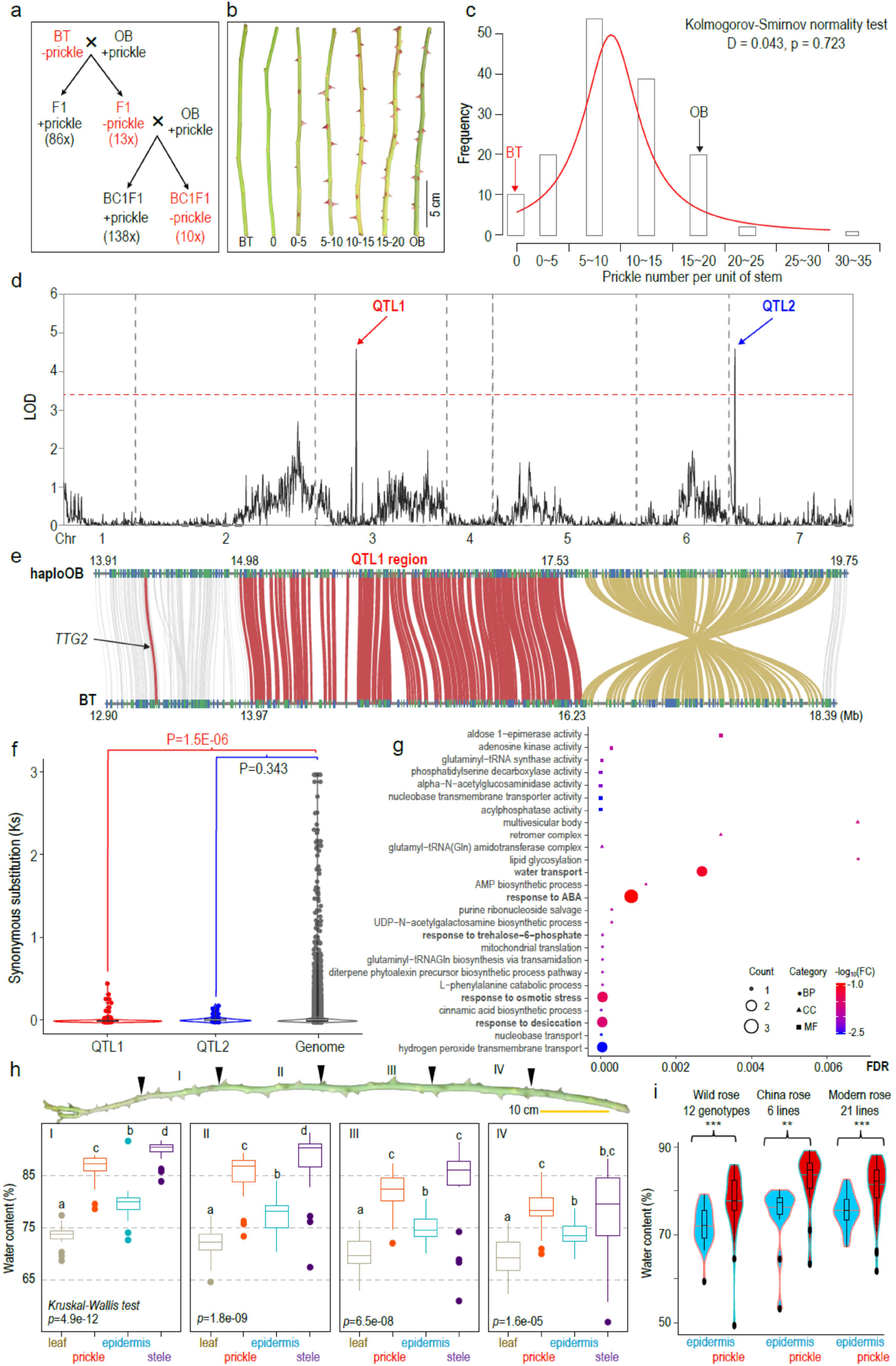
Genetic analysis of stem prickle inheritance in OB x BT populations. **a.** Inheritance of prickle distribution on stems of F1 and BC1F1 populations. Plants without prickles on stems indicated with a ‘-’ in red color while a ‘+’ labeled plants have prickles. Numbers in brackets mark the plant number featuring each trait category. **b.** Typical phenotypes of prickle density on stems of the parents (BT and OB) and the BC1F1 population. Numbers indicate the range (0, >0~5, >5~10, >10~15, >15~20; the same for **c**) of prickle density per 20cm unit of stem. **c.** Normal distribution of stem prickle density in BC1F1 population. The X-axis indicates prickle density per unit of stem length. Y-axis marks the frequency of BC1F1 individuals showing the range of prickles. Arrows show prickle features of both parents (BT in red and OB in dark). **d.** QTL analysis for prickle density on shoot in BC1F1 population using multiple QTL mapping (MQM) method. The red line indicates the significance threshold (=3.4). Red and blue arrows mark the two QTL over threshold on Chr3 and Chr7, respectively. **e.** Syntenic gene pairs (in red lines) between OB and BT surrounding the QTL1. This QTL was neighboring a genome conversion (brown lines) between OB and BT. Potential *TTG2* labeled with a dark red line outside of QTL1. Numbers on/under chromosomes show physical positions in Mb. **f.** Comparison of synonymous substitutions (*Ks*) between the QTL regions and the rest of the genome. **g.** GO enrichment pattern for differentially expressed genes located in the two QTL. X-axis shows FDR values; FC, enrichment fold change against genome level; BP, biological process; CC, cellular components; MF, molecular function. GO terms in bold relate to water usage. **h.** Relative water contents are significantly higher in prickles (red) than in epidermis (blue), steles (dark purple) or leaves (brown). Data for one modern rose genotype shown. I, II, III, and IV represent every three nodes marked by dark arrow heads. Letters (a, b, c, and d) above each boxplot indicate pairwise significances (p<0.01; non-parametric *Wilcoxon test*) for eight biological replicates. **i.** Prickles (red) with relatively higher water contents than epidermis (blue; p<0.001, *Student’s t-test*) in a collection of randomly selected roses (12 wild rose, 6 China rose, and 21 modern cultivated genotypes). ***, P<0.001; **, P<0.01. For h and i, prickle number does not co-vary with water content for either prickle, epidermis or stele.

### Stem prickle density is regulated by two main QTL

We conducted a QTL analysis using the 2,160 SNP markers for the BC1F1 population [25]. Three and two QTL were identified using interval mapping (IM; Fig. S24) and multiple QTL mapping (MQM; Fig. 3d) methods, respectively. The QTL1 (the logarithm of the odds ratio, LOD=4.59; 12.3% variance explained) and QTL2 (LOD=4.58; 12.4% variance) were identical (LOD threshold 3.4). An additive-effect was identified for OB alleles in both QTL1 (3.98) and QTL2 (4.23). QTL1 was located on Chr3 next to a genome inversion event (Fig. 3e). This 2.26 Mb QTL (13.97-16.23 Mb) harbored 153 genes in BT and covered a 2.55 Mb fragment (14.98-17.53 Mb; 254 genes) in the haploOB genome. QTL1 was close to, but did not co-locate with, the QTL regions regulating continuous flowering, double flower, or self-incompatibility traits (Table S29) [14]. However, this QTL was indeed within the ~15 Mb segment linked with prickle density on stems as reported previously by Hibrand Saint-Oyant et al [14] and Zhou et al [32] (Table S30). QTL1 had 117 syntenic genes between BT and haploOB. The *TTG2-like* (Rw3G013440/ Chr3G0468221), a candidate proposed previously [14], was located ~0.6 Mb outside of QTL1 vicinity. QTL2 was on Chr7 with 60 syntenic genes between BT (BT, 19.03-20.28 Mb, 73 genes) and OB (haploOB, 21.23-22.34 Mb, 65 genes; Fig. 3d; Fig. S25). The weak QTL3 was located to the beginning of Chr7 (BT, 0.98-2.64 Mb, 204 genes; haploOB, 1.08-2.73 Mb, 211 genes; 177 syntenic, Fig. S26, S27). QTL2 and QTL3 were not identified previously in genetic analyses with F1 populations [14, 32], suggesting that populations in different generations might provide different powers in genetic analyses.

Genes in the QTL1 and QTL3 regions showed a significant reduction of the synonymous-substitutions (*Ks*) compared to the rest of the genome (Fig. 3f; Fig. S27; Data S1, Table S31). Additionally, the genes in the QTL1 displayed a lower level of non-synonymous to synonymous substitution ratios (*Ka/Ks*) compared to the rest of Chr3 (Fig. S28; Table S31), implying a purifying selection. A detailed examination of the syntenic genes in the QTL regions showed that genes involved in water usage and desiccation responses were significantly and differentially expressed in the young shoot tips (<0.5 cm in length) between BT and OB (Fig. 3g; Fig. S29). Two genes encoding for aquaporin PIP2s, known factors involved in coordination of cell growth and drought adaptation [33, 34], featured substantial expression reduction in shoot tips of OB than in BT (Fig. S30). Our results raised a possibility that genes for prickle patterning might hitchhike on the sequences related to physiological adaptations for water accumulation and storage, or vice versa. Prickles may have a relatively high water-content.

We measured the relative water content in prickles, epidermis, steles, and leaves at different sections of the stems of two genotypes (*R.* ‘Cinderella’, and *R.* ‘Tianshan Xiangyun’; Fig. 3h; Fig. S31-S34). Prickles always had higher water content than epidermis and leaves in the four sections of seasonal shoots. In contrast, relative water content dropped sharply in prickles on older shoots, while maintaining similar levels in leaves compared to young shoots (Fig. S32; Table S32). This modifies the original hypothesis. Prickles may serve as water storage structures at early phases graduating to defense mechanisms with stem maturation. Prickles might have different functions over the life span of these woody plants. A further examination of water contents in prickles and epidermis for a collection of 39 lines (12 wild, 6 China roses, and 21 modern genotypes) revealed a similar pattern (Fig. 3i; Fig. S35, S36; Table S33). Taken conjointly, prickle patterning may hitchhike on the “guild” of genes involved in water maintenance in rose stems [18]. Consequently, prickles remain absent in natural populations of *R. wichuraiana* as shrubs grow in moist, mesophytic habitats with monsoon climates [17, 18]. Additionally, a preliminary phylogenetic analysis with chloroplast genome sequences indicated independent origins of prickle-free roses (Table S34; Fig. S37).

### Expression variation of candidate genes in QTL and trichome GRN associates with stem prickle patterning

Rose prickles are deformed plant trichomes [16] and the responsible GRN was well studied in *Arabidopsis* [1, 5, 35–38]. Our scanning-electronic-microscopy (SEM) analyses revealed that, in contrast to the normal initiation of prickles and stomata development on OB shoots, BT maintained the ability to develop stomata but lacked prickle initiation (Fig. 4a). Therefore, we identified potential candidate genes involved in the trichome GRN, and focused on the MBW-complex, i.e., the MYB-, bHLH- and WRKY-like transcription factors in the QTL regions [1, 5, 35, 37, 38]. In *Arabidopsis*, this MBW complex involves the physical interaction between GL3 with GL1 and TTG1, respectively, to initiate trichome formation. *TTG2* acts genetically downstream of this complex while sharing partial activity with *GL2* to initiate trichome development [36]. A complete loss of *TTG2* and *GL2* activity caused the failure of trichome initiation [39]. Single repeat R3-MYBs act as negative regulators of trichome development via competition with the R2R3-type MYBs to form a repressor complex.

**Fig. 4.**
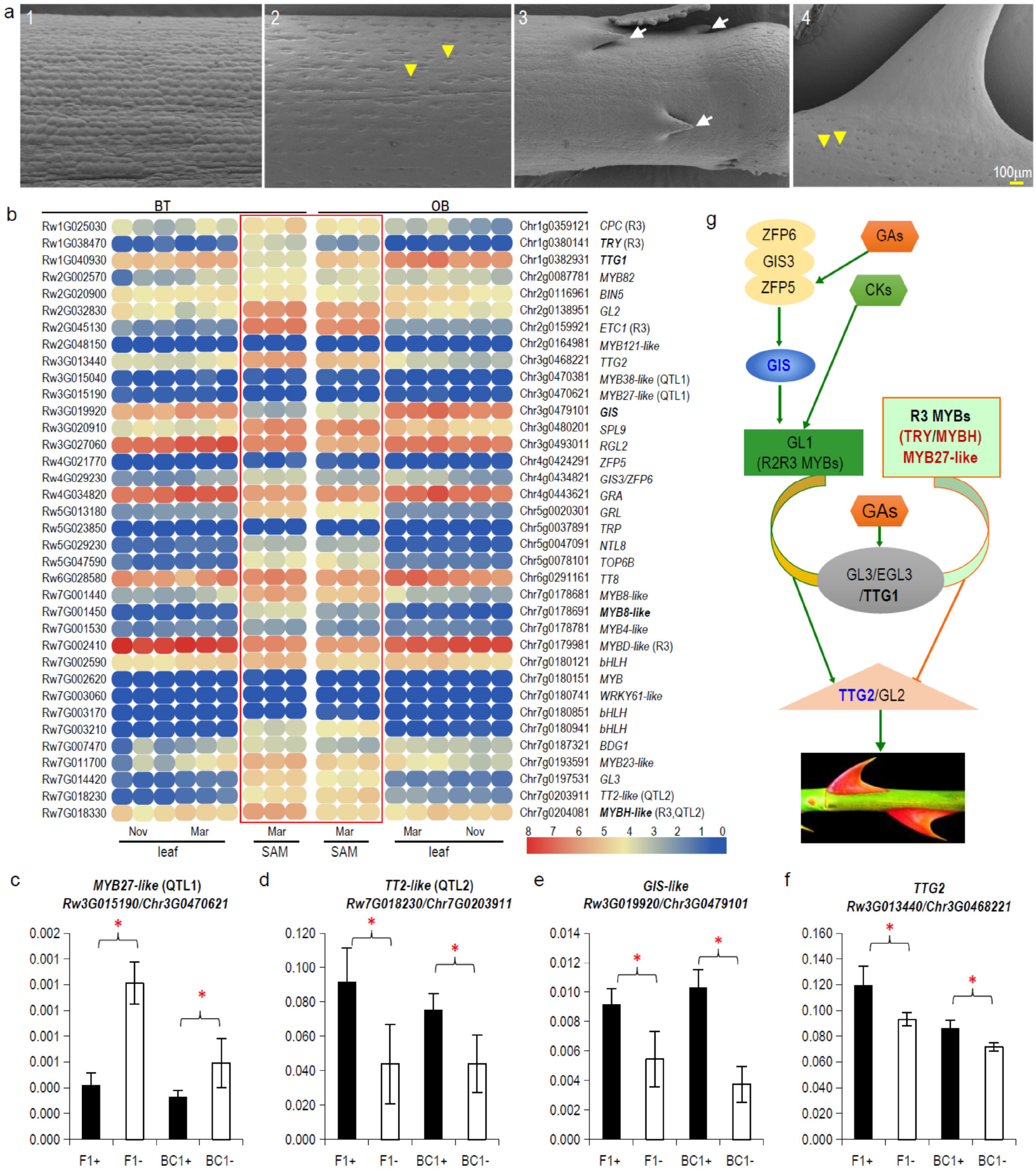
Identification of candidate genes involved in stem prickle patterning. **a.** SEM analysis of early (**1** and **3**) and older (**2** and **4**) stems for BT (**1** and **2**) and OB (**3** and **4**). Prickles and stomata are marked with white arrows and yellow arrow heads, respectively. **b** Expression heatmap of genes involved in prickle-initiation GRN. The bar at the right bottom marked the normalized FPKM values for both BT and OB (red denoted high expression and blue indicated low expression). Names in bold mark genes showing minimum two-fold changes at p<0.05 (*Student’s two tailed t-test*). R3 type MYBs indicated with “R3” in brackets. Genes in QTL marked in brackets. SAM Mar (in red box), young shoots in March; leaf Mar/Nov, with expanding leaves in March/November. **c-f**. Expression comparisons for candidate genes (*MYB27-like*, **c**; *TT2-like*, **d**; *GIS-like*, **e**; *TTG2-like*, **f**) in F1 and BC1F1 pools with (F1+ or BC1+) or without (F1- or BC1-) prickles on stem. **g**. A simplified GRN regulating rose prickle initiation based on information from *Arabidopsis*.

*Arabidopsis* has ~150 trichome-related genes involved in initiation, development, birefringence and other processes (Data S2). We identified 240 BT homologous genes with ten and one being CPG (OB gain compared to BT; 1,314 genes) and CPL (OB lost; 193 genes), respectively. Among these, the OB-lost but BT-featuring *Rw2G037200* (a duplication of *Rw2G037100*) encoded potentially a TT2-like protein with the At5G35550, which encoded for TT2, as the closet homolog in Arabidopsis [40]. However, both genes were expressed at very low level in both BT and OB (Fig. S38). The remainings were related to trichome branching (Data S2). None of these 11 genes was in the vicinity of QTL region. We next focused primarily on those 29 genes known to regulate trichome initiation and development (Fig. 4b; Data S3, S4). In comparison to trichome GRNs of *Arabidopsis*, cucumber and cotton, we detected no presence-absence polymorphisms of genes between BT and OB. These results indicate that the trichome GRN is highly conserved among these roses [3, 5, 6, 41].

We found four *MYB*s (*Rw3G015040/ Chr3g0470381*, *Rw3G015190/Chr3g0470621* for QTL1; *Rw7G018230/ Chr7g0203911* and *Rw7G018330/Chr7g0204081* for QTL2). We identified five *MYB*s (including a homolog of *WER/GL1/MYB23-like*), three *bHLH*s, and one *WRKY* in QTL3 (Data S3, S4; Table S35; Fig. S38-S43). We detected no significant sequence variation in the means of *Ka*/*Ks* change between these orthologous genes. We found no lineage-specific expansion/contraction of these three gene families. The interpretation here is that the regulation of gene expression, not the encoding potential, of these trichome-related genes may underlie regulation of prickle-free stems in BT. Indeed, we detected five genes with at least two-fold significant expression changes in very young shoots including apical meristems between BT and OB (<0.5cm; Fig. 4b) [20]. The expression of positive regulators *GIS* and *TTG1* decreased, while the negative regulators R3-type *MYB*s (*TRY-* and *MYBH-like*) and *MYB8-like* increased in BT compared to OB (Fig. 4b, Data S4). A qRT-PCR assay confirmed the significant expression variation of *MYB27-like* in QTL1 (Rw3G015190/Chr3G0470621), *TT2-like* in QTL2 (Rw7G018230 /Chr7G0203911), *GIS-like* (Rw3G019920/Chr3G0479101) and *TTG2-like* in both F1 and BC1F1 pools with or without prickles (Fig. 4c-f). Both MYB27-like and TT2-like TFs are proposed to regulate anthocyanin synthesis in *Petunia* and *Arabidopsis* via modulating the MBW-complex [42–44]. However, their functional roles in trichome development need further investigation. Acting downstream of the R3-MYBs, GIS promoted trichome development via transcriptional regulation of genes involved in MBW-complex [45, 46]. Featuring strong sequence variations in the upstream transcriptional regulatory regions (Fig. S45), the WRKY *TTG2* worked downstream of MBW-complex but partially independent of *GL2.* This leads to the regulation of trichome patterning and seed-coat anthocyanin biosynthesis [36, 39]. It was a candidate for stem prickle patterning in roses [14]. Interestingly, variations in *cis-*regulatory mutations in MYB TFs, *TCL* and *TRY*, together with a *GL1*mutation, triggerred the trichome development on *Arabidopsis* fruits. These variations were significantly associated with low spring precipitation thus may contribute to climate adaptation [47]. Taken together, we propose that gene expression variation of these candidates underlies prickle-development variation between BT and OB (Fig. 4g).

## Conclusions

We generated a chromosome-level genome assembly for the heterozygous diploid BT, and identified the complex inheritance pattern of stem prickles. The production of the diploid BT genome sequence provides us with a foundation and novel resources to study rose biology and mine molecular markers associated with important traits. They offer targeted strategies for breeding new rose varieties and for studying the genomes evolution in the commercially important family, Rosaceae. For long-term benefits, according to the fairy tales of Jacob L.K. Grimm & Wilhelm C. Grimm (1812, *Little Brier-Rose*) and Oscar Wilde (1888, *The Nightingale and the Rose*) respectively, landscapes of prickle-free roses should reduce fatalities in populations of male royalty and birds in the genus *Luscinia.*

## Methods and Materials

The detailed descriptions of methods are available as Supplementary Materials at *NSR* online.

## Availability of supporting data

All data supporting the results of this study are included in the manuscript and its additional files. Genome assembly and annotations were deposited in the NCBI BioProject under accessions PRJNA542225.

## Ethics statement

Not applicable.

## Author contributions

J-Y.H. conceptualized the project; J-Y.H., D-Z.L. and K.T. coordinated the research. X.J., W.W., Y.S., D.W., W.C., and Z.S. collected the samples, extracted the genomic DNA and total RNA and carried out the prickle phenotyping. X.J. did the qRT-PCR. M.Z., X.D., G.Y., X.C. and X.L. analyzed and visualized the data. J-Y.H. wrote the paper. All authors have read and approved the final manuscript.

## Acknowledgements

We thank Prof. David H. Byrne for BT materials. We appreciate Shubin Li, Shulan Chen, Hongyuan Yu and Yuanlin Lv for plants cultivation, Zhenhua Guo, Haitao Cui, Hong Wang, Peter Bernhardt and Peter Raven for discussions. This work is partially facilitated by the Germplasm Bank of Wild Species, iFlora HPC Center of GBOWS, and the Kunming Botanical Garden, KIB, CAS.

